# Optogenetic inhibition reveals distinct contributions of medial prefrontal cortex to intertemporal choice in young and aged rats

**DOI:** 10.1101/2025.07.15.664949

**Authors:** Mojdeh Faraji, Caesar M. Hernandez, Alexa-Rae Wheeler, Todd J. Sahagian, Scott W. Harden, Charles J. Frazier, Matthew R. Burns, Barry Setlow, Jennifer L. Bizon

**Author notes:** These authors contributed equally to this work. **Correspondance:** Jennifer L. Bizon, Ph.D. Department of Neuroscience University of Florida PO Box 100244 Gainesville, FL 32610-0244.

## Abstract

The ability to choose adaptively between rewards differing in magnitude and delay (intertemporal choice) is critical for numerous life outcomes. Compared to younger adults, older adults tend to exhibit greater preference for large, delayed over small, immediate rewards (i.e., less delay discounting), which could lead to missed opportunities to obtain resources necessary for quality of life. Intertemporal choice is mediated by the prefrontal cortex, but how this is impacted by advanced age is not well understood. We used optogenetic inactivation to investigate contributions of medial prefrontal cortex (mPFC) during distinct components of an intertemporal choice task in young and aged rats. mPFC inactivation during deliberation (during decisions between small, immediate vs. large, delayed rewards) increased preference for large, delayed rewards in both age groups. In contrast, inactivation during delays prior to large reward delivery increased preference for large, delayed rewards only in aged rats. Choices were unaffected by inactivation during other task phases. Results suggest that mPFC integrates information regarding anticipated outcomes into the decision process across the whole lifespan, but that only in aging is mPFC critical for consolidating information regarding reward delays into the decision structure in order to modulate choice behavior.

---

Intertemporal choice refers to decisions between smaller, more immediate rewards and larger rewards that are more delayed in their arrival. Longer delays to arrival of the larger rewards reduce their subjective value, causing an increase in preference for smaller, more immediate rewards (i.e., delays discount the value of the larger rewards – (Ainslie 2005, Odum 2011)). Alterations in intertemporal choice are evident in numerous neuropsychiatric conditions, including substance use disorders, anorexia, schizophrenia, obsessive compulsive disorder, ADHD, and apathy, and can contribute substantially to declines in measures of quality of life and activities of daily living (Beauchaine, Ben-David et al. 2017, Steinglass, Lempert et al. 2017, Brown, Hart et al. 2018, Amlung, Marsden et al. 2019, Ong, Graves et al. 2019, Sloan, Sanches et al. 2023, Brown, Sofis et al. 2024). Even in the absence of pathology, patterns of decision making during intertemporal choices can change across the lifespan, with greater preference for smaller, immediate rewards (i.e., greater “impulsive choice”) in childhood and adolescence shifting in young adulthood toward increasing preference for larger, more delayed rewards, with (at least in some cases) an even further shift toward delayed gratification in advanced ages (Green, Fry et al. 1994, Scheres, Dijkstra et al. 2006, Jimura, Myerson et al. 2011, Eppinger, Nystrom et al. 2012, Buono, Whiting et al. 2015, Halfmann, Hedgcock et al. 2016), although see (Seaman, Abiodun et al. 2022). Greater preference for delayed over immediate gratification such as can occur with age may be advantageous in certain circumstances; however, such changes can also be maladaptive, resulting in disadvantageous decisions regarding health care and finances. Complicating matters further, aging is a major risk factor for neurodegenerative conditions such as Alzheimer’s and Parkinson’s diseases, which are associated with their own shifts in intertemporal choice (Gleichgerrcht, Ibáñez et al. 2010, Lebreton, Bertoux et al. 2013, Bayard, Jacus et al. 2014, Thoma, Maercker et al. 2017, Hernandez, McCuiston et al. 2024). As such, it is important to understand how this form of decision making is regulated across the lifespan.

Intertemporal choice is regulated by neural networks that include the striatum, amygdala, and prefrontal cortex (PFC), with strong evidence suggesting that in humans, the PFC plays a particularly important role (Peters and Büchel 2011, Samanez-Larkin and Knutson 2015, Bailey, Simpson et al. 2016, Fobbs and Mizumori 2017). Findings from rodent (particularly rat) models are consistent with this conclusion. Expression of genes involved in neurotransmitter signaling in the medial PFC (mPFC) is associated with variability in intertemporal choice in young adult rats, and both pharmacological and genetic manipulations of mPFC in young adult rats cause shifts in decision making in intertemporal choice tasks (Churchwell, Morris et al. 2009, Loos, Pattij et al. 2010, Feja and Koch 2014, Freund, MacGillivilray et al. 2014, Sonntag, Brenhouse et al. 2014, Meda, Freund et al. 2019, Wenzel, Zlebnik et al. 2023). Importantly, as in humans, aging in rats is accompanied by reductions in impulsive choice, consistent with age-related changes in other aspects of PFC-mediated cognition in this species (Barense, Fox et al. 2002, Simon, LaSarge et al. 2010, Bizon, Foster et al. 2012, Roesch, Esber et al. 2012, Beas, Setlow et al. 2013, Hernandez, Vetere et al. 2017, Tomm, Tse et al. 2018, Cammarata and De Rosa 2022). In addition, previous work from our labs has linked age-dependent attenuation in GABA(B) receptor expression and receptor activity in the prelimbic subregion of the mPFC to greater preference for large, delayed over small, immediate rewards (Hernandez, McQuail et al. 2022), providing the start of a mechanistic understanding of age changes in intertemporal choice.

Biochemical approaches and genetic or pharmacological manipulations have enabled determination of specific receptor and molecular substrates of intertemporal choice within mPFC; however, these approaches lack the temporal specificity to enable definition of the specific components of the decision process (e.g., deliberation vs. outcome evaluation) in which mPFC is involved. Approaches such as electrophysiology and functional neuroimaging provide such temporal specificity, but they are limited in their capacity to draw conclusions regarding causal roles for the neural activity recorded. We and others have recently employed optogenetics to begin to draw causal inferences regarding the temporally-precise contributions of activity in specific brain regions and circuits during cost-benefit decision making (Zalocusky, Ramakrishnan et al. 2016, Orsini, Hernandez et al. 2017, Bercovici, Princz-Lebel et al. 2018, Bercovici, Princz-Lebel et al. 2023, McLaughlin and Redish 2023, Truckenbrod, Betzhold et al. 2023, White, Morningstar et al. 2024). These optogenetic studies indicate critical roles for encoding of different types of information in these structures across the decision process. A further study that employed this approach in young adult and aged rats revealed age-specific contributions of BLA to processing information during distinct components of intertemporal choice (Hernandez, Orsini et al. 2019). Here we extend this work to investigate the role of mPFC in distinct components of intertemporal choices in young adulthood and aging.

## Materials and Methods

### Subjects

Young adult (6 months old) and aged (24 months old) Fischer 344 x Brown Norway F1 hybrid (FBN) rats were obtained from the National Institute on Aging colony (Charles River Laboratories) and individually housed in the Association for Assessment and Accreditation of Laboratory Animal Care International-accredited vivarium facility in the McKnight Brain Institute building at the University of Florida. Because these studies were conducted at a time at which aged female FBN rats were largely unavailable from the National Institute on Aging, only male rats were used. The vivarium facility was maintained at a constant temperature of 25°C with a 12-hour light/dark cycle (lights on at 0700) and free access to food and water except as otherwise noted. Rats were acclimated in this facility and handled for at least one week prior to initiation of any procedures. All animal procedures were conducted in accordance with the rules and regulations of the University of Florida Institutional Animal Care and Use Committee and National Institutes of Health guidelines. A subset of rats completed testing in only some of the behavioral epochs due to illness or mishap, and some rats were excluded entirely for misplaced injections. In addition, some of the rats were trained in a delayed response working memory task prior to testing in the intertemporal choice task. Only the final numbers of rats included in each analysis are provided below.

### Surgical Procedures

Surgical procedures were performed as in our previous work (Orsini, Hernandez et al. 2017). Rats were anesthetized with isofluorane gas (1-5% in O_2_) and secured in a stereotaxic frame (David Kopf). An incision along the midline over the skull was made and the skin was retracted. Bilateral burr holes were drilled above the mPFC and four additional burr holes were drilled to fit stainless steel anchoring screws. Bilateral guide cannulae (22-gauge, Plastics One) were implanted to target the mPFC (anteroposterior: +2.7 mm from bregma, mediolateral: ±0.7 mm from bregma, dorsoventral: 1.5 mm from the skull surface) and secured to the skull using dental cement. A total of 0.7 µL per hemisphere of a 3.5 × 10^12^ vg/ml titer solution (University of North Carolina Vector Core) containing AAV5 packaged with either halorhodopsin (CamKIIα-eNpHR3.0-mCherry, n=13 young and n=8 aged rats) or mCherry alone (CamKIIα-mCherry, n=5 young and n=5 aged rats) was delivered via an injection needle inserted into the implanted cannulae over 3 minutes (at a dorsoventral coordinate 2 mm below the cannula tips). Stainless steel obdurators were placed into the cannulae to minimize the risk of infection. Immediately after surgery, rats received subcutaneous injections of meloxicam (2 mg/kg). Meloxicam was also administered 24, 48, and 72 hours post-operation. A topical ointment was applied as needed to facilitate wound healing. Sutures were removed after 10-14 days and rats recovered for at least 2 weeks before food restriction and behavioral testing began.

### Ex Vivo Electrophysiology

For *ex vivo* electrophysiological verification of halorhodopsin function, young adult (n=2) and aged (n=3) rats underwent surgery as described above except that neither guide cannulae nor skull screws were implanted (Hernandez, Orsini et al. 2019). Three weeks after surgery, rats were administered a ketamine/xylazine cocktail (100 mg/kg ketamine, 10 mg/kg xylazine, i.p.) and adequate anesthesia was evaluated by the lack of response to hind paw pinch. Rats were then transcardially perfused with an ice-cold sucrose laden artificial cerebrospinal (ACSF) solution containing (in mM): 205 sucrose, 10 dextrose, 1 MgSO4, 2 KCl, 1.25 NaH2PO4, 1 CaCl2, 25 NaHCO3, oxygenated with carbogen (95% O2, 5% CO2). Brains were extracted, submerged in sucrose laden ACSF, and sectioned coronally to prepare 300-micron thick slices through the PFC using a vibratome (Leica VT1200S). Slices were transferred to a low-calcium high-magnesium ACSF containing (in mM): 124 NaCl, 10 dextrose, 3 MgSO4, 2.5 KCl, 1.23 NaH2PO4, 1 CaCl2, 25 NaHCO3, oxygenated with carbogen, and maintained at 35 °C for 30 minutes. Slices were then permitted to cool to room temperature passively for at least 30 minutes, followed by transfer to a recording chamber (Warner Instruments RC-26GLP) perfused at 2 mL/min with standard ACSF containing (in mM): 126 NaCl, 11 dextrose, 1.5 MgSO4, 3 KCl, 1.2 NaH2PO4, 2.4 CaCl2, 25 NaHCO3, oxygenated with carbogen, and maintained at 30°C using an inline heater (Warner Instruments SH-28 and TC-324C). Patch pipettes were pulled from borosilicate glass (Sutter Instrument BF150-86-10) using a Flaming/Brown micropipette puller (Sutter Instrument P-97) and had an open-tip resistance of 4-6 MΩ. Patch pipettes were filled with a potassium gluconate-based internal solution containing (in mM): 125 K-gluconate, 10 phosphocreatine, 1 MgCl2, 10 HEPES, 0.1 EGTA, 2 Na2ATP, 0.25 Na3GTP, adjusted to pH 7.3 by adding KOH, and adjusted to 290 mOsm by adding H2O. Data are presented uncorrected for the liquid junction potential. Patch clamp experiments were performed using an upright stereomicroscope (Olympus BX-WI) under DIC-IR optics. Fluorescence imaging of mCherry and activation of halorhodopsin was achieved using a TTL-controlled light source (Excelitas technologies, X-Cite 110LED) paired with a green excitation / red emission filter cube (Omega Optical XF414). Electrophysiological recordings were performed using Clampex 10.7, a Multiclamp 700B patch-clamp amplifier with a CV-7B headstage, and a DigiData 1440A digitizer (Molecular Devices). Voltage clamp data were recorded at 20 kHz and low pass filtered using a 4-pole Bessel filter with a 2 kHz cutoff frequency. Offline analysis was performed in OriginPro using custom software written by CJF.

### Behavioral Testing Procedures

#### Apparatus

Testing was conducted in 12 identical standard rat behavioral test chambers (Coulbourn Instruments) with metal front and back walls, transparent Plexiglas side walls, and a floor composed of steel rods (0.4 cm in diameter) spaced 1.1 cm apart. Each test chamber was housed in a sound-attenuating cubicle and was equipped with a custom food pellet delivery trough fitted with a photobeam to detect head entries (TAMIC Instruments), located 2 cm above the floor and extending 3 cm into the chamber in the center of the front wall. A nosepoke hole equipped with a 1.12 W lamp for illumination was located directly above the food trough. Food rewards consisted of 45 mg purified ingredient rodent food pellets (Test Diet; 1811155 5TUM). Two retractable levers were positioned to the left and right of the food trough (11 cm above the floor). A 1.12 W house light was mounted near the top of the rear wall of the sound-attenuating cubicle. For optogenetics, laser light (561 nm, 8–10 mW output at the fiber tip, Shanghai Laser & Optics Century) was delivered through a patch cord (200 µm core, Thor Labs) to a rotary joint (1 X 2, 200 µm core, Doric Lenses) mounted above the operant chamber. At the rotary joint, the light was split into 2 outputs. Tethers (200 µm core, 0.39 NA, Thor Labs) connected these outputs to the optic fibers implanted in the mPFC. A computer running Graphic State 4.0 software (Coulbourn Instruments) was used to control the behavioral apparatus and laser delivery, and to collect data.

#### Behavioral shaping and initial training

The intertemporal choice task was based on a design by (Evenden and Ryan 1996) and was used previously to demonstrate age-related alterations in intertemporal choice in both Fischer 344 (Simon, LaSarge et al. 2010) and FBN (Hernandez, Vetere et al. 2017, Hernandez, Orsini et al. 2019, Hernandez, McQuail et al. 2022) rats. Prior to commencement of behavioral testing, both young and aged rats were food-restricted to 85% of their free-feeding weights over the course of 5 days and maintained at these weights for the duration of the experiments. Rats were initially shaped to lever press to initiate delivery of a food pellet into the food trough and were then trained to nosepoke to initiate lever extension. Each nosepoke during this stage of shaping initiated extension of either the left or right lever (randomized across pairs of trials), a press on which yielded a single food pellet. After two consecutive sessions of reaching criterion performance (45 presses on each lever), rats began testing on the intertemporal choice task.

#### Intertemporal choice task

Task procedures were modified as in our previous work (Hernandez, Orsini et al. 2019). Briefly, each 60-min session consisted of 3 blocks of 20 trials each (Figure 1a). The trials were 60 s in duration and began with a 10 s illumination of both the nosepoke port and house light. Entry into the nosepoke port during this time extinguished the nosepoke light and triggered lever extension. Any trials on which rats failed to nosepoke during this 10 s window were scored as omissions. Each 20-trial block began with 4 forced choice trials, in which either the right or left lever was extended, in order to remind rats of the delay contingencies in effect for that block. These forced choice trials were followed by 16 free choice trials, in which both levers were extended. For all trials, one lever (either left or right, counterbalanced across age groups) was always associated with immediate delivery of one food pellet (the small reward), and the other lever was always associated with 3 food pellets (the large reward) delivered after a variable delay. Lever assignments to small vs. large reward remained consistent throughout testing. Within a session, the duration of the delay preceding large reward delivery increased across the three blocks of trials. The actual delay durations were adjusted individually for each rat, such that the percent choice of the large reward corresponded as closely as possible to 100% in block 1, 66% in block 2, and 33% in block 3. On all trials, rats were given 5 s to press a lever, after which the levers were retracted and food was delivered into the food trough (either immediately or after the programmed delay). If rats failed to press a lever within 5 s, the levers were retracted, lights were extinguished, and the trial was scored as an omission. An inter-trial interval (ITI) followed either food delivery or an omitted trial, after which the next trial began.

**Figure 1.**
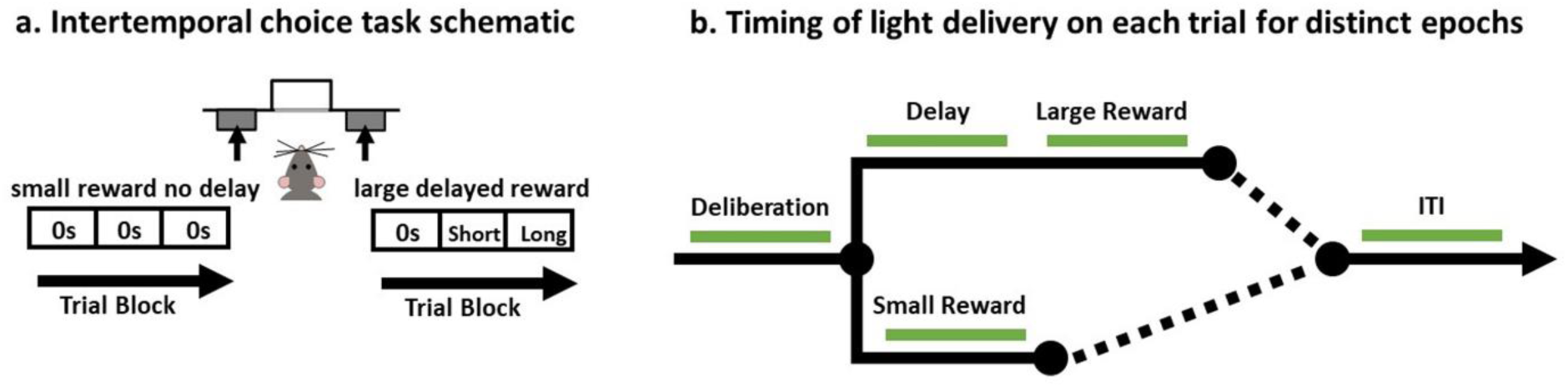
Schematics of intertemporal choice task and timing of light delivery. **a)** Schematic of the intertemporal choice task illustrating the choices and trial blocks across which the duration of the delay to the large reward increased. On each trial, rats were presented with two response levers that differed with respect to the magnitude and timing of associated reward delivery. Presses on one lever delivered a small (one food pellet), immediate reward, whereas presses on the other lever delivered a large (three food pellets) reward delivered after a variable delay. Trials were presented in a blocked design, such that the delay to the large reward increased across successive blocks of trials in a session. **b)** Schematic of a single trial in the intertemporal choice task showing the task epochs during which light was delivered (represented by the green line). Using a within-subjects design, light was delivered during *deliberation* (when levers are presented until a choice is made); *small reward delivery*; *delay*; *large reward delivery*; and *intertrial interval (ITI)*.

Rats were initially trained for 15 sessions on the intertemporal choice task. They were then lightly anesthetized and optic fibers (200 µm core, 0.39 NA; Precision Fiber Products) were placed into the guide cannulae such that they extended 1.5 mm past the end of the guide cannulae and were cemented in place. After recovery, rats resumed training but were now tethered to the rotary joint.

#### Evaluation of optogenetic inactivation during specific task epochs

The effects of temporally-discrete optogenetic inhibition of mPFC were tested in both young and aged rats using a within-subjects design (Figure 1b). Data from three consecutive sessions occurring just prior to inactivation sessions (in which rats did not receive light delivery) were used as the baseline against which to compare the effects of mPFC inactivation. Note that inactivation took place only on free-choice trials. Task epochs in which the mPFC was inactivated were as follows: *deliberation* (light delivery began 500 ms prior to illumination of the nosepoke light and continued until the rat pressed the lever, for a maximum of 15 s); *small reward delivery* (light delivery began when food was dispensed and remained on for 6 s); *large reward delivery* (light delivery began when food was dispensed and remained on for 6 s), *delay* (light delivery began upon press of the large reward lever and remained on throughout the delay intervals ranging between 1-50 s); *intertrial interval (ITI*; light delivery began 14 seconds after reward was dispensed and continued for 4 s). Finally, the sequence of mPFC inactivation sessions during each task epoch was counterbalanced across rats.

#### Virus Expression and Cannula Placement Histology

After completion of behavioral testing, rats were administered a lethal dose of Euthasol (sodium pentobarbital and phenytoin solution; Virbac, Fort Worth, TX, USA) and perfused transcardially with a 4°C solution of 0.1M phosphate buffered saline (PBS), followed by 4% (w/v) paraformaldehyde in 0.1M PBS. Brains were removed and post-fixed for 24 h, then transferred to a 20% (w/v) sucrose solution in 0.1M PBS for 72 h (all chemicals purchased from Fisher Scientific, Hampton, NH, USA). Brains were sectioned at 35 µm using a cryostat maintained at -20°C. Sections were rinsed in 0.1M TBS and incubated in blocking solution consisting of 3% normal donkey serum, 0.3% Triton-X-100 in 0.1M TBS for 1 h at room temperature. Sections were then incubated with rabbit anti-mCherry antibody (ab167453, Abcam, Cambridge, MA, USA) diluted in blocking solution at a dilution of 1:1000 (72 hours, 4°C). Following primary incubation, sections were rinsed in 0.1M TBS and incubated in blocking solution containing the secondary antibody (donkey anti-rabbit conjugated to AlexaFluor-488, 1:300) for 2 h at room temperature. After rinsing in 0.1M TBS, sections were mounted on electrostatic glass slides and coverslipped using Prolong Gold (Thermo Fisher Scientific, Waltham, MA, USA). Sections were visualized at 20X using an Axio Imager 2 microscopy system (Carl Zeiss Microscopy, LLC, Thornwood, NY, USA) to assess mCherry expression in mPFC neurons. Cannula placements and mCherry expression were mapped onto plates adapted from the rat brain atlas of Paxinos and Watson (2005).

### Experimental Design and Statistical Analysis

#### Evaluation of age differences in halorhodopsin effects on mPFC neuronal activity

Data analysis was performed using OriginLab and custom electrophysiology analysis software written by CJF. Electrophysiological measures of intrinsic properties and mean firing were compared between young and aged cells, or with vs. without activation of halorhodopsin, using unpaired Student’s t-tests. Welch’s correction for unequal variance was applied when necessary.

#### Evaluation of mPFC inactivation on intertemporal choice

The effects of light delivery were tested separately for each task epoch (*deliberation, small reward delivery, large reward delivery, delay,* and *ITI*). For each epoch, comparisons were made using a mixed factor ANOVA (laser condition × age × delay block), with age (2 levels: young and aged) as a between-subjects factor, and laser condition (2 levels: laser on or off) and delay block (3 levels: zero, short delay, long delay) as within-subjects factors. For all analyses, the Greenhouse-Geisser correction was used to account for violations of sphericity in ANOVAs. To better understand significant main effects or interactions, further analyses were conducted separately in each age group using a two-factor ANOVA (laser condition × block). Note that for those epochs in which effects of mPFC inactivation during the delay were tested, data analyses were confined to blocks 2 and 3, as there was no delay in block 1. Alpha was set to 0.05 for all statistical analyses. For statistical results that reached the threshold for significance, η_p_^2^ (partial eta squared) was used to report the effect size for mixed-factor ANOVAs, and 1-β was used to report the observed power.

#### Evaluation of choice strategy resulting from mPFC inactivation

Additional analyses were conducted to better understand the shifts in choice performance following mPFC inactivation during the deliberation, delay, and large reward epochs. Trials were categorized based on choices made on the previous trial. For each epoch, trials were categorized as “small-<u>shift</u>-to-large” or “large-<u>stay</u>-on-large”. The number of “small-<u>shift</u>-to-large” trials was divided by the total number of “small reward choice” trials, and the number of “large-<u>stay</u>-on-large” trials was divided by the total number of “large reward choice” trials in that session and expressed as a percentage. For each task epoch assessed, percentages of trials in each category were compared using a mixed factor ANOVA, with age as a between-subjects factor and laser condition as a within-subjects factor.

#### Effects on other task performance measures resulting from mPFC inactivation

Other task measures were compared between mPFC inactivation and baseline conditions in task epochs in which mPFC inactivation produced significant changes in choice behavior. Specifically, on free choice trials, response latency (the time between lever extension and a lever press) was compared. Previous work shows that response latencies can differ for large and small reward levers (Hernandez, Vetere et al. 2017), and hence analyses were conducted separately for each lever using data from delay block 2, during which rats made roughly equivalent numbers of choices on each reward lever. Response latency and total number of trials completed were compared using a mixed factor ANOVA (age × laser condition).

## Results

### Electrophysiological confirmation of light-induced inhibition of mPFC neurons expressing eNpHR3.0

To assess the functional effects of halorhodopsin activation, mPFC neurons expressing mCherry were evaluated in young and aged rats using patch-clamp technique. Pyramidal neurons were identified by their location, morphology, and electrophysiological properties (Table 1). In halorhodopsin-expressing neurons from young and aged rats, green light produced an outward current in voltage-clamp configuration and hyperpolarized neurons from rest in current-clamp mode (Table 1, Figure 2a). Current steps were used to depolarize neurons above action potential threshold to achieve continuous firing, and exposure to green light reduced action potential firing rates in both age groups (Figure 2b). Specifically, in response to intermediate stimuli of 200 to 400 pA, activation of halorhodopsin reduced mean firing frequency observed in young PFC neurons from 13.78 ± 0.7 Hz to 4.44 ± 0.99 Hz (t=7.71, p<0.001). In PFC neurons from aged animals, activation of halorhodopsin similarly reduced the mean action potential firing rate observed in response to these stimuli from 16.23 ± 1.15 Hz to 8.24 ± 1.36 Hz (t=4.49, p<0.001). This represents a reduction in mean firing frequency observed of 9.34 ± 0.90 Hz in young neurons and of 7.99 ± 1.62 Hz in aged neurons, which was not statistically different (t=0.73, p=0.47). These findings confirm that the strategies used to deliver opsins are successful and demonstrate opsin functionality: green light hyperpolarizes halorhodopsin-expressing neurons, and effectively reduces neuronal activity in response to excitatory stimuli in both young and aged animals.

**Figure 2.**
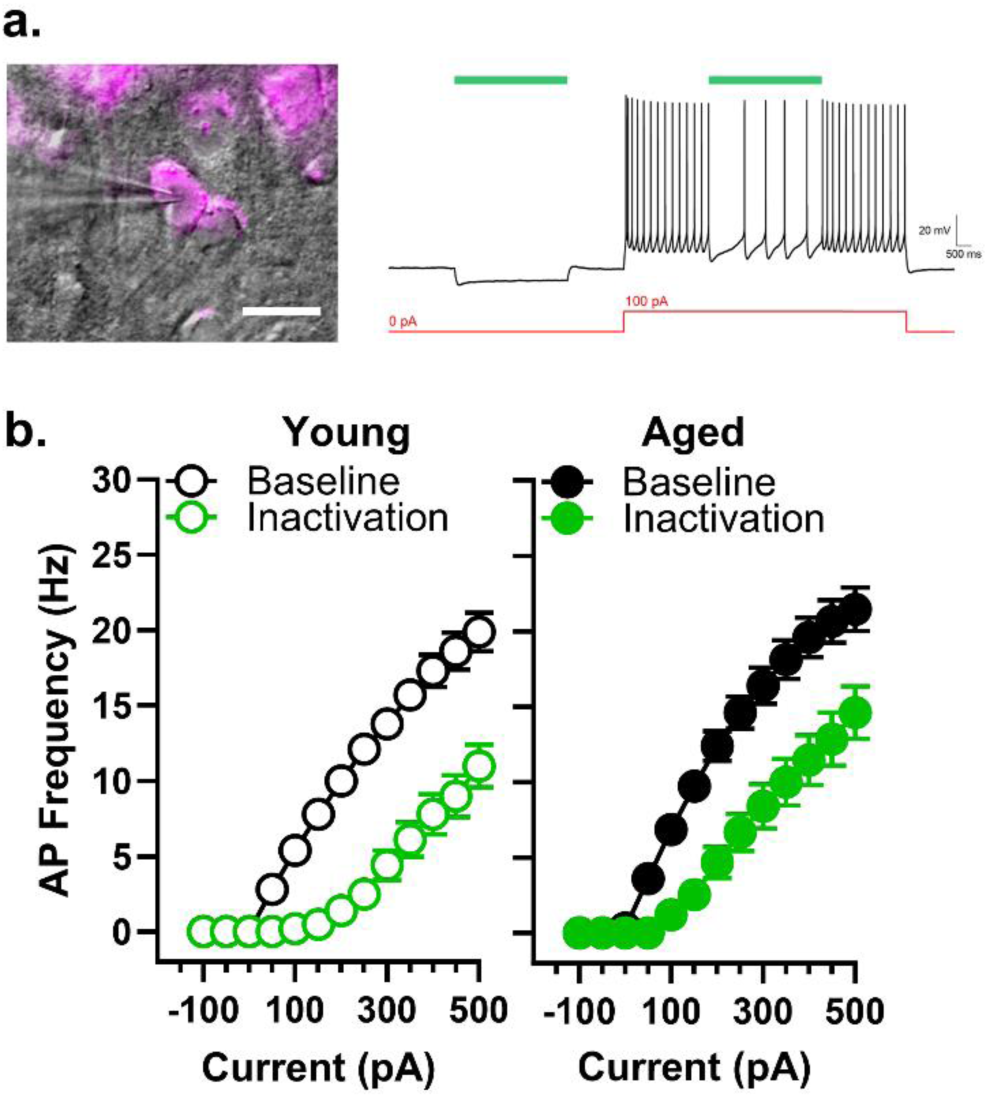
Functional inhibition of mPFC pyramidal neurons via activation of halorhodopsin. **a)** left: mPFC pyramidal neurons expressing mCherry were identified by their morphology and electrophysiological properties; right: representative current-clamp recording demonstrating that halorhodopsin activation by green light (green bar) hyperpolarizes neurons at rest (left) and reduces firing rate in cells that continuously fire in response to excitatory current injection (right). **b)** Halorhodopsin-expressing neurons from both young and aged animals showed reduced neuronal gain (reduced firing rate in response to direct current injection) when halorhodopsin is concurrently activated by exposure to green light.

**Table 1.** Electrophysiological Characteristics of mCherry-Expressing mPFC Neurons.

| Metric | Halo Young | Halo Aged |  |
| --- | --- | --- | --- |
| Number of Neurons Tested | 10 | 15 |  |
| Holding Current at -70 mV (pA) | -82.86 ±9.03 | -90.28 ±20.85 | t=0.33, p=0.75 |
| Membrane Resistance (MΩ) | 154.06 ±10.90 | 143.83 ±18.95 | t=0.47, p =0.64 |
| Whole-Cell Capacitance (pF) | 159.89 ±10.63 | 145.29 ±9.23 | t=1.03, p=0.31 |
| Resting Membrane Potential (mV) | -55.28 ±1.76 | -56.47 ±2.28 | t=0.38, p=0.71 |
| Maximum Firing Frequency (Hz) | 23.85 ±1.15 | 25.13 ±1.81 | t=0.53, p=0.62 |
| Opsin Current (pA) | 263.32 ±29.70 | 191.13 ±44.06 | t=1.22, p=0.24 |
| Opsin Hyperpolarization (mV) | -29.33 ±2.50 | -21.64 ±3.91 | t=1.47, p=0.15 |

### Fiber placement and AAV transduction

Expression of mCherry was used to confirm viral transduction in the mPFC of rats used in behavioral studies that were injected with either AAV5-CamKIIα-eNpHR3.0-mCherry (AAV-eNpHR3.0, white circles in Figure 3) or AA5-CamKIIα-mCherry alone (AAV-control, black circles in Figure 3). Cannula placements were centered in the dorsal mPFC, and the brain volumes transduced by AAV-eNpHR3.0 and AAV-control (calculated from the atlas of Paxinos & Watson, 2005) were comparable in young and aged rats.

**Figure 3.**
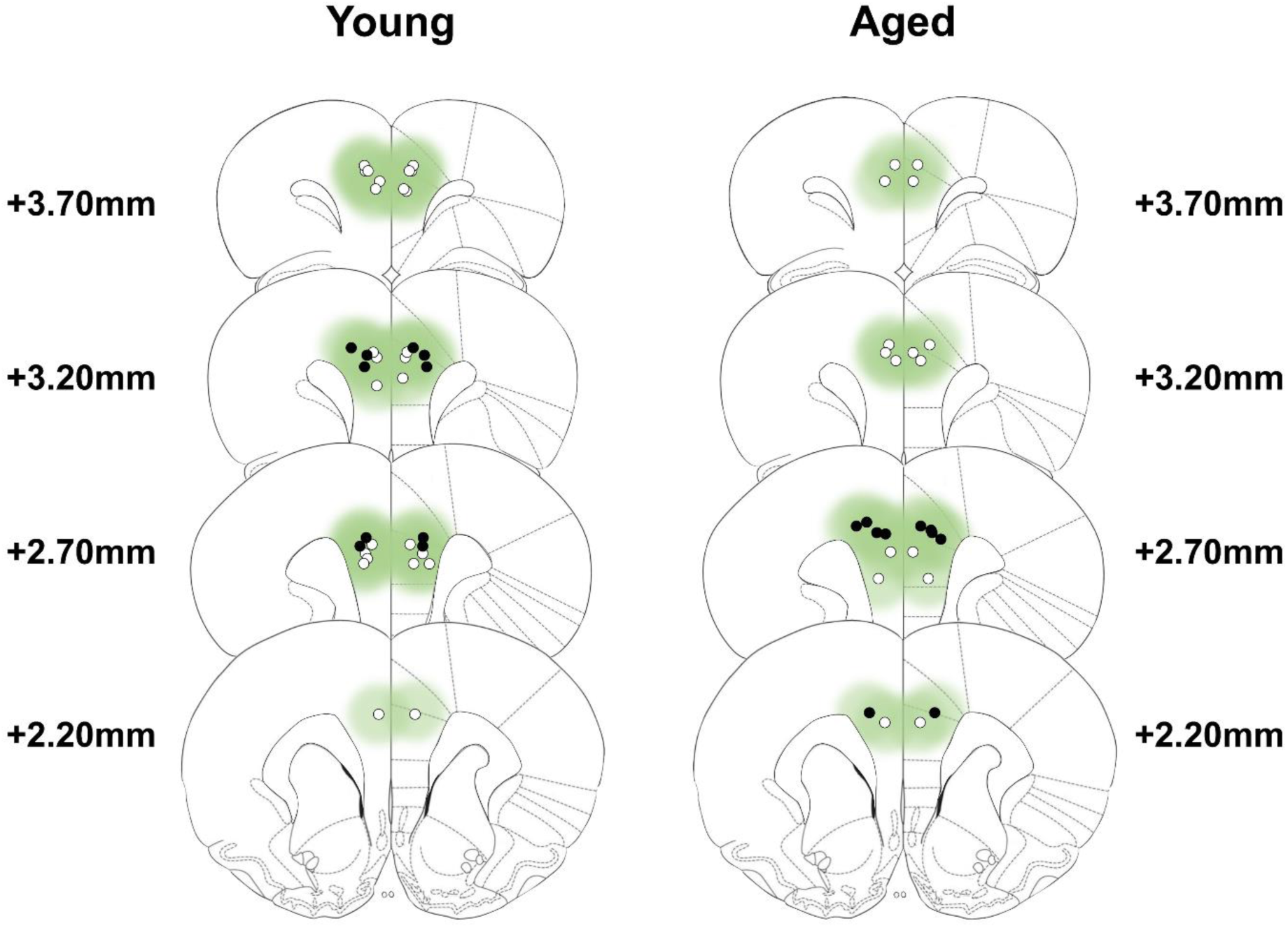
Verification of viral expression and fiber optic placements. Viral-transduction in young (left) and aged (right) rats is depicted in green. Green color intensity indicates the extent of expression of AAV5-CamKIIα-eNpHR3.0-mCherry or AA5-CamKIIα-mCherry. Open circles represent optic fiber placements in the experimental (eNpHR3.0) groups, and filled black circles represent optic fiber placements in the control groups. Viral expression and fiber placements are mapped to standardized coronal sections corresponding to +3.70 mm through +1.70 mm from bregma according to the atlas of Paxinos and Watson (2006).

### Effects on choice behavior of mPFC inactivation during the deliberation epoch

Inactivation of the mPFC during the deliberation epoch (n=13 young, n=8 aged) increased choice of the large reward in young and aged rats (Figure 4a). A three-factor ANOVA (laser condition × age × delay block) indicated a main effect of laser condition (F_(1,19)_=7.08, p=0.02, η_p_^2^=0.27, 1-β=0.71) such that rats chose the large reward more frequently when mPFC was inactivated in comparison to baseline. There was an expected main effect of delay block (F_(1.85,35.07)_=136.19, p<0.01, η_p_^2^=0.88, 1-β=1.00) such that rats chose the large reward less frequently as the delay associated with the large reward increased. The interaction between laser condition and delay block did not reach statistical significance (F_(1.80,34.17)_=3.06, p=0.07), and no main effect of age (F_(1,19)_=0.06, p=0.81), age × laser condition interaction (F_(1,19)_=0.30, p=0.59), age x delay interaction (F_(1.85,35.07)_=0.09, p=0.76), or age × laser condition × delay interaction (F_(1.80,34.17)_=0.24, p=0.77) were observed.

**Figure 4.**
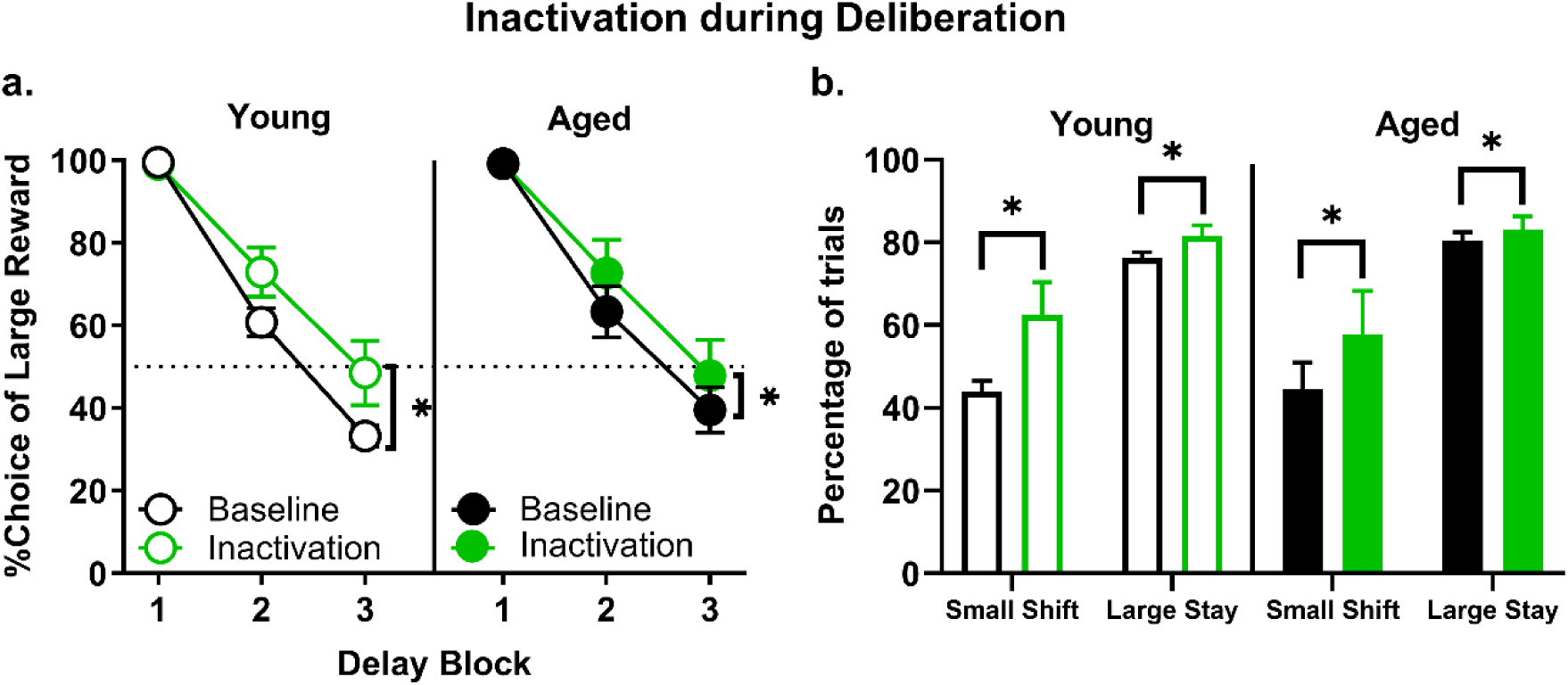
Effect of mPFC inactivation during the deliberation epoch. **a)** Inactivation of the mPFC during the deliberation epoch (prior to a choice) resulted in an increase in preference for the large, delayed reward across both young (n = 13) and aged (n = 8) rats. **b)** Analysis of trial-by-trial choice strategies revealed that the increased choice of the large, delayed reward caused by mPFC inactivation during deliberation (panel a) was due to both an increase in the percentage of trials on which rats shifted to the large, delayed reward following a choice of the small, immediate reward and an increase in the percentage of trials on which rats stayed on the large, delayed reward following a choice of the large, delayed reward. Error bars represent standard error of the mean (SEM). * = main effect of laser.

The data above show that, irrespective of age, mPFC inactivation during the deliberation epoch altered choice behavior by increasing preference for the large reward. Trial-by-trial analyses were subsequently conducted to determine the effects of mPFC inactivation on two distinct behavioral strategies that could mediate these shifts in choice preference. Specifically, this analysis determined the degree to which mPFC inactivation influenced rats to “shift” to the large reward following a choice of the small reward on the previous trial, versus “stay” with the large reward option following choice of the large reward on the previous trial. As shown in Figure 4b, the percentage of trials during deliberation epoch inactivation on which a large reward choice was followed by a second large reward choice (large-stay) increased as a function of laser condition (main effect of laser condition: F_(1,19)_=4.81, p=0.04, η_p_^2^=0.20, 1-β=0.55), but there were no main effects or interactions involving age (main effect of age: F_(1,19)_=0.91, p=0.35; laser condition × age: F_(1,19)_=0.54, p=0.47). A similar analysis of the percentage of trials on which a choice of the small reward was followed by choice of the large reward (small-shift) revealed that mPFC inactivation increased the percentage of such shifts (main effect of laser condition: F_(1,19)_=5.05, p=0.04, η_p_^2^=0.21, 1-β=0.57), but again, this effect was independent of age (main effect of age: F_(1,19)_=0.09, p=0.77, laser condition × age interaction: F_(1,19)_=0.16, p=0.70).

Other task measures were compared between mPFC inactivation and baseline conditions using a two-factor ANOVA (age x laser condition), with age as the between-subjects factor and laser condition as the within-subjects factor. As shown in Table 2, the number of trials completed in a session decreased as a function of laser condition (F_(1,19)_=7.13, p=0.02, η_p_^2^=0.27, 1-β=0.72), but no main effect or interaction associated with age was present. Table 3 shows latencies to press the small and large reward lever on free-choice trials. Neither mPFC inactivation during the deliberation epoch nor age affected latencies to press the large reward lever. Latencies to press the small lever were longer when mPFC was inactivated during the deliberation epoch (F_(1,16)_=5.36, p=0.03, η_p_^2^=0.25, 1-β=0.57), but this measure was not affected by age.

**Table 2.** Effects of mPFC inactivation on number of trials completed per session.

| Epoch | Age | Condition | Mean | SEM | Statistical Comparisons |
| --- | --- | --- | --- | --- | --- |
| Deliberation | Young | Baseline | 45.67 | 1.01 | **Laser Condition:** $F_{(1,19)}=7.13$, $p=0.02$<br>**Age:** $F_{(1,19)}=0.65$, $p=0.43$<br>**Laser Condition × Age:** $F_{(1,19)}=0.15$, $p=0.71$ |
|  |  | Inactivation | 43.08 | 2.04 |  |
|  | Aged | Baseline | 44.21 | 1.29 |  |
|  |  | Inactivation | 40.75 | 2.60 |  |
| Delay | Young | Baseline | 29.14 | 2.68 | **Laser Condition:** $F_{(1,17)}=3.26$, $p=0.09$<br>**Age:** $F_{(1,17)}=1.61$, $p=0.22$<br>**Laser Condition × Age:** $F_{(1,17)}=2.36$, $p=0.14$ |
|  |  | Inactivation | 28.92 | 2.45 |  |
|  | Aged | Baseline | 27.91 | 3.97 |  |
|  |  | Inactivation | 25.14 | 4.06 |  |
| Outcome<br>(large reward) | Young | Baseline | 46.53 | 3.75 | **Laser Condition:** $F_{(1,16)}=3.26$, $p=0.09$<br>**Age:** $F_{(1,16)}=4.19$, $p=0.06$<br>**Laser Condition × Age:** $F_{(1,16)}=1.59$, $p=0.23$ |
|  |  | Inactivation | 46.08 | 3.77 |  |
|  | Aged | Baseline | 44.5 | 8.14 |  |
|  |  | Inactivation | 42 | 8.59 |  |

**Table 3.**
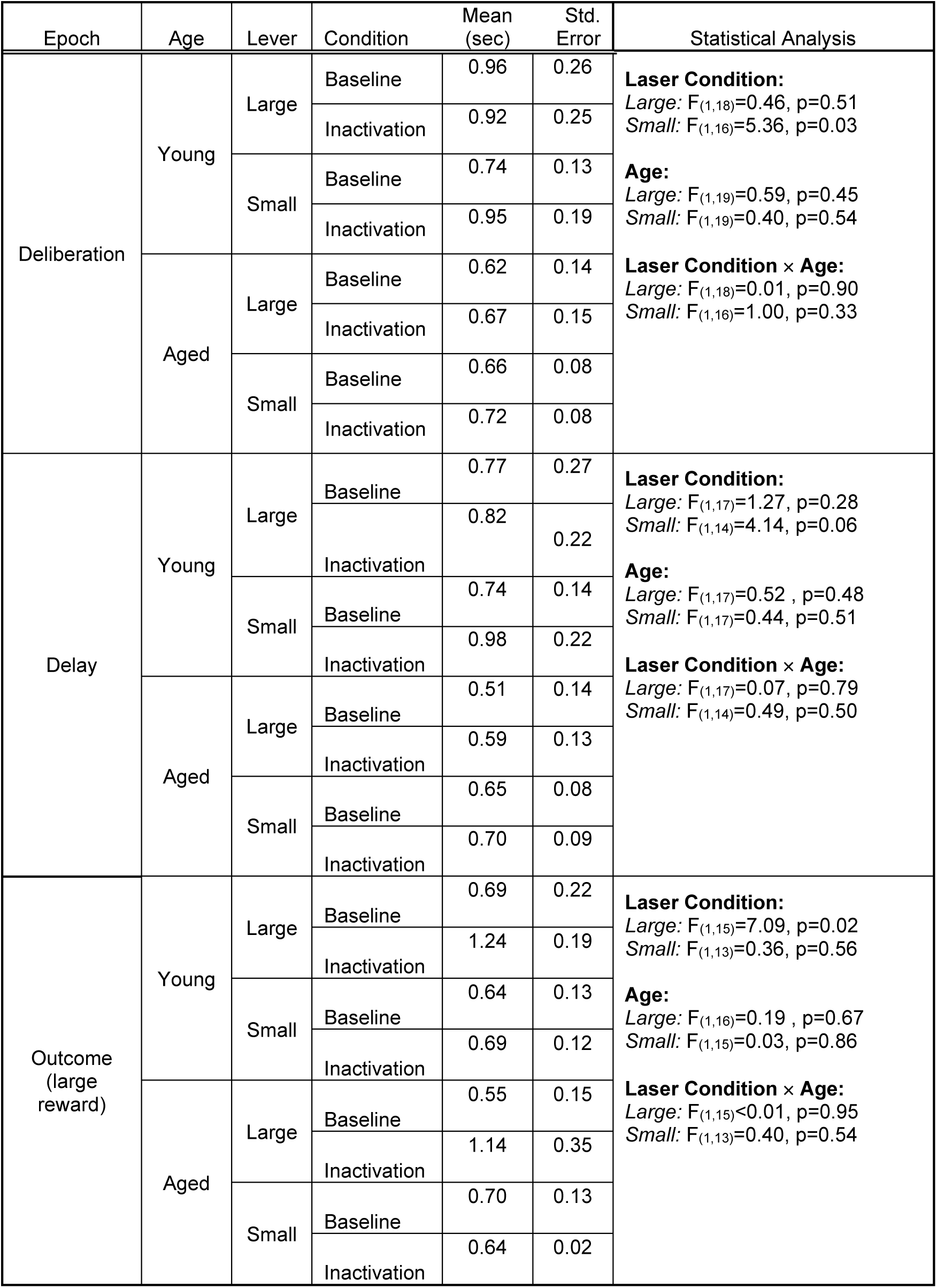
Effects of mPFC inactivation on lever response latencies in free choice trials.

#### Effects of mPFC inactivation during the delay epoch

The effects of mPFC inactivation during the delay epoch (n=12 young and n=7 aged) were tested in delay blocks 2 and 3 using a three-factor ANOVA (laser condition × age × delay block). As expected, there was a main effect of delay block (F_(1,17)_=37.08, p<0.01, η_p_^2^=0.69, 1-β=1.00) such that both young and aged rats decreased their choice of the large reward as the delay prior to large reward delivery increased (Figure 5a). Importantly, significant differences in choice behavior resulted from mPFC inactivation during the delay epoch (main effect of laser condition: F_(1,17)_=14.72, p<0.01, η_p_^2^=0.46, 1-β=0.95) such that rats increased their choice of the large reward when mPFC was inactivated, and this effect was larger in aged rats compared to young (laser condition × age: F_(1,17)_=5.68, p=0.03, η_p_^2^=0.25, 1-β=0.61). There was no main effect of age (F_(1,17)_=1.74, p=0.21), nor were there interactions associated with delay (laser condition × delay block: F_(1,17)_=0.75, p=0.40; age × delay block: F_(1,17)_=0.32, p=0.58; laser condition × age × delay block: F_(1,17)_ = 0.10, p=0.76). To further investigate the nature of the observed mPFC inactivation effects, follow-up two-factor ANOVAs (laser condition × delay block) were performed in each age group. Consistent with the significant interaction observed in the omnibus ANOVA, these analyses revealed that mPFC inactivation significantly increased choice of the large reward in aged rats (main effect of laser condition: F_(1,6)_=14.60, p<0.01, η_p_^2^=0.71, 1-β=0.89, main effect of delay block: F_(1,6)_=7.79, p=0.03, η_p_^2^=0.57, 1-β=0.65; laser condition × delay block: F_(1,6)_=0.33, p=0.58), but not in young rats (main effect of laser condition: F_(1,11)_=1.47, p=0.25; main effect of delay block: F_(1,11)_=42.95, p<0.01, η_p_^2^=0.80, 1-β=1.00; laser condition × delay block: F_(1,11)_=0.32, p=0.59).

**Figure 5.**
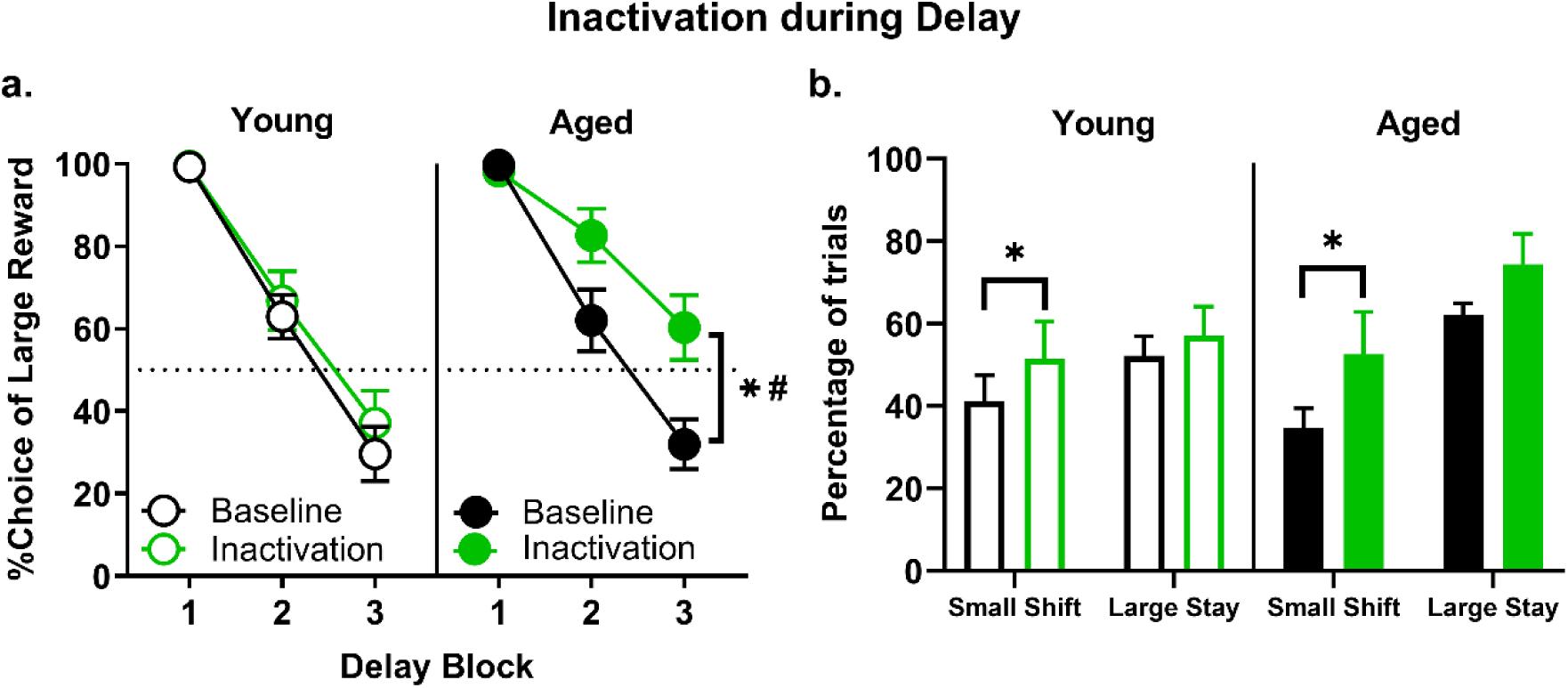
Effect of mPFC inactivation during the delay epoch. **a)** Inactivation of the mPFC during the delay epoch resulted in a significant increase in preference for the large, delayed reward in aged (n = 7), but not young (n = 12) rats. **b)** Analysis of trial-by-trial choice strategies in blocks 2 and 3 revealed that the percentage of trials on which rats shifted to the large, delayed reward following a choice of the small, immediate reward (small-shift) increased as a result of mPFC inactivation during delay epoch. Error bars represent standard error of the mean (SEM). * = main effect of laser; # = age x laser interaction.

A trial-by-trial analysis was conducted on sessions in which inactivation took place during the delay epoch (Figure 5b). mPFC inactivation increased the percentage of trials in blocks 2 and 3 on which a small reward choice was followed by choice of the large reward (small-shift) (main effect of laser condition: F_(1,17)_=6.05, p=0.03, η_p_^2^=0.26, 1-β=0.64) although there was no main effect or interaction involving age (main effect of age: F_(1,17)_=0.07, p=0.80; laser condition × age interaction: F_(1,17)_=0.42, p=0.53). Similarly, the percentage of trials during delay epoch inactivation on which a large reward choice was followed by a second large reward choice (large-stay) showed a trend toward an increase as a function of laser condition (main effect of laser condition: F_(1,17)_=3.84, p=0.07), but there was no main effect or interaction involving age (main effect of age: F_(1,17)_=2.94, p=0.10; laser condition × age: F_(1,17)_=0.73, p=0.41).

Other task measures were compared between mPFC inactivation and baseline conditions following delay epoch inactivation using a two-factor ANOVA (age x laser condition). The number of trials completed (Table 2) was not affected (main effect of laser condition: F_(1,17)_=3.26, p=0.09), nor did this measure differ between age groups (main effect of age: F_(1,17)_=1.61, p=0.22; laser condition x age: F_(1,17)_=2.36, p=0.14). As depicted in Table 3, neither mPFC inactivation during the delay epoch nor age affected latencies to press either lever.

### Effects of mPFC inactivation during the large reward epoch

The effects of inactivation of the mPFC during delivery of the large reward (n=12 young and n=6 aged) were tested using a three-factor ANOVA (laser condition × age × delay block; Figure 6a). This analysis revealed the expected main effect of delay block (F_(1.84,29.49)_= 104.12, p<0.01, η_p_^2^=0.87, 1-β=1.00) such that both young and aged rats decreased their choice of the large reward as the delay to large reward delivery increased, but no main effects of laser (F_(1,16)_=1.41, p=0.25) or age (F_(1,16)_=1.25, p=0.28), nor interactions between laser condition and delay block (F_(1.96,31.30)_=0.96, p=0.39), age and delay block (F_(1.84,29.49)_=0.93, p=0.40), or age and laser condition (F_(1,16)_=1.37, p=0.26). There was, however, a trend toward an interaction between laser condition, age, and delay block (F_(1.96,31.30)_=2.86, p=0.07) such that aged but not young rats chose the large reward more frequently at the longest delay (block 3) when mPFC was inactivated.

**Figure 6.**
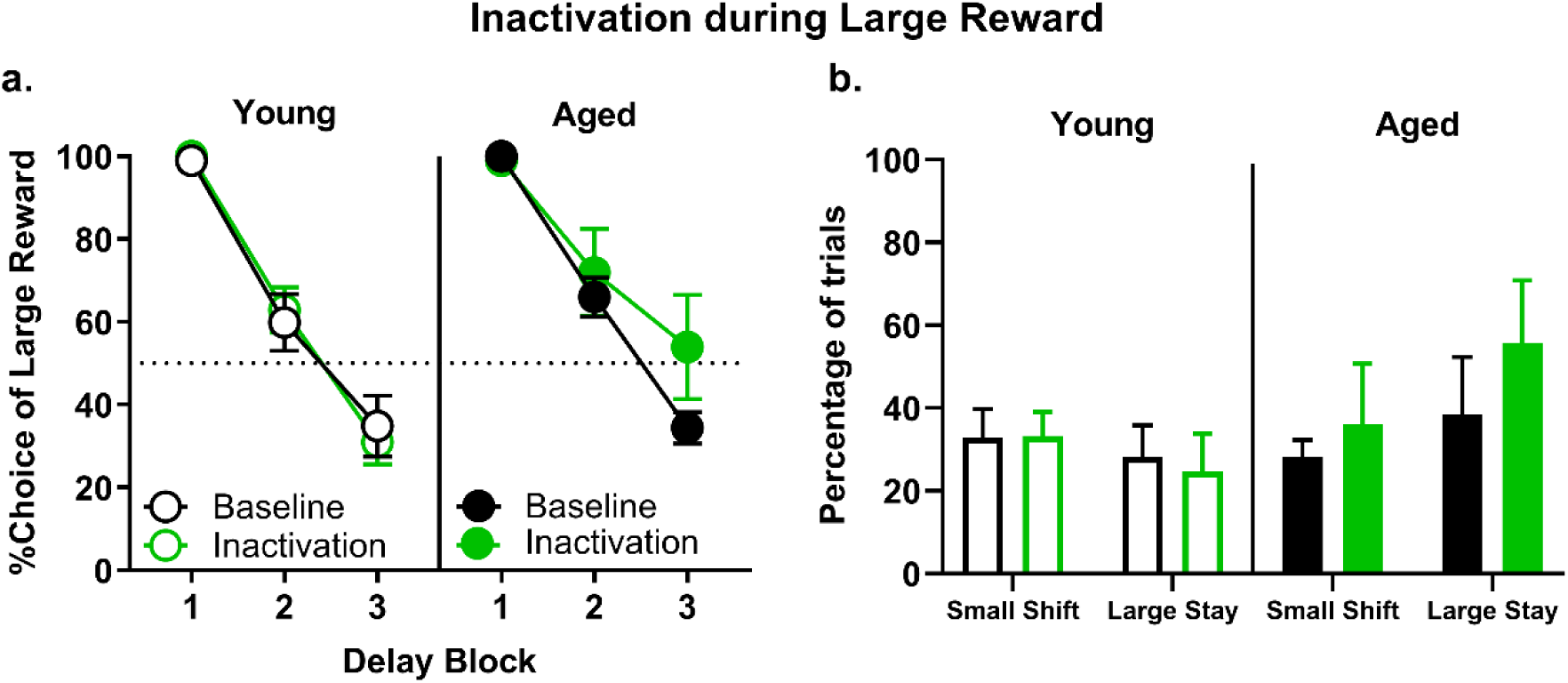
Effects of mPFC inactivation during the large reward epoch. **a)** Inactivation of mPFC during the large reward epoch resulted in a numerical but not statistically significant increase in preference for the large reward in aged rats at the longest delay (aged, n = 6; young, n = 12). **b)** Analysis of trial-by-trial choice strategies in block 3 revealed no effect of mPFC inactivation during large reward delivery on the percentage of trials in which aged rats stayed on the large, delayed reward following a choice of the large, delayed reward (large-stay), or the percentage of trials on which rats shifted to the large, delayed reward following a choice of the small, immediate reward (small-shift). Error bars represent standard error of the mean (SEM).

Because mPFC inactivation during the large reward epoch produced a striking numerical increase in choice of the large reward in aged but not young rats in the long-delay block (block 3, Figure 6a), an analysis of choice strategy specifically at long delays (block 3) was conducted (Figure 6b). The results showed a numerical increase in the percentage of trials in which a large reward choice was followed by a second large reward choice (large-stay) in aged rats when mPFC was inactivated, but the effect did not reach statistical significance (F_(1,15)_=0.87, p=0.37); There was also no main effect of age (F_(1,16)_=2.30, p=0.15) nor an interaction between age and laser condition (F_(1,15)_=1.70, p=0.21). The percentage of trials in block 3 on which a small reward choice was followed by choice of the large reward (small-shift) also did not differ by age (F_(1,16)_=0.01, p=0.92), or laser condition (F_(1,15)_=0.32, p=0.58), nor was there any age x laser condition interaction (F_(1,15)_=0.28, p=0.60).

On free choice trials, the number of trials completed/session (Table 2) was not affected by mPFC inactivation during large reward delivery (main effect of laser condition: F_(1,16)_=3.26, p=0.09; main effect of age: F_(1,16)_=4.19, p=0.06; age x laser condition: F_(1,16)_=1.59, p=0.23). In contrast, mPFC inactivation led to longer latencies to press the large reward lever (main effect of laser condition: F_(1,15)_=7.19, p=0.02, η_p_^2^=0.33, 1-β=0.71). See Tables 2 and 3 for full statistical results of these analyses.

### Effects on choice behavior of mPFC inactivation during small reward delivery

mPFC inactivation during the small reward epoch (n=12 young and n=6 aged) only showed the expected main effect of delay block (F_(1.90,30.44)_=109.62, p<0.01, η_p_^2^=0.87, 1-β=1.00) such that both young and aged rats decreased their choice of the large reward as the delay prior to the large reward increased (Figure 7a). A three-factor ANOVA (laser condition × age × delay block) indicated no main effects of laser condition (F_(1,16)_=0.15, p=0.71), or age (F_(1,16)_=0.04, p=0.86), nor were any interaction effects observed (laser condition × delay block: F_(1.87,29.97)_= 0.10, p = 0.90; laser condition × age: F_(1,16)_=0.20, p=0.66; age × delay block: F_(1.90,30.44)_=0.16, p= 0.86; laser condition × age × delay block: F_(1.87,29.97)_=0.13, p=0.87).

**Figure 7.**
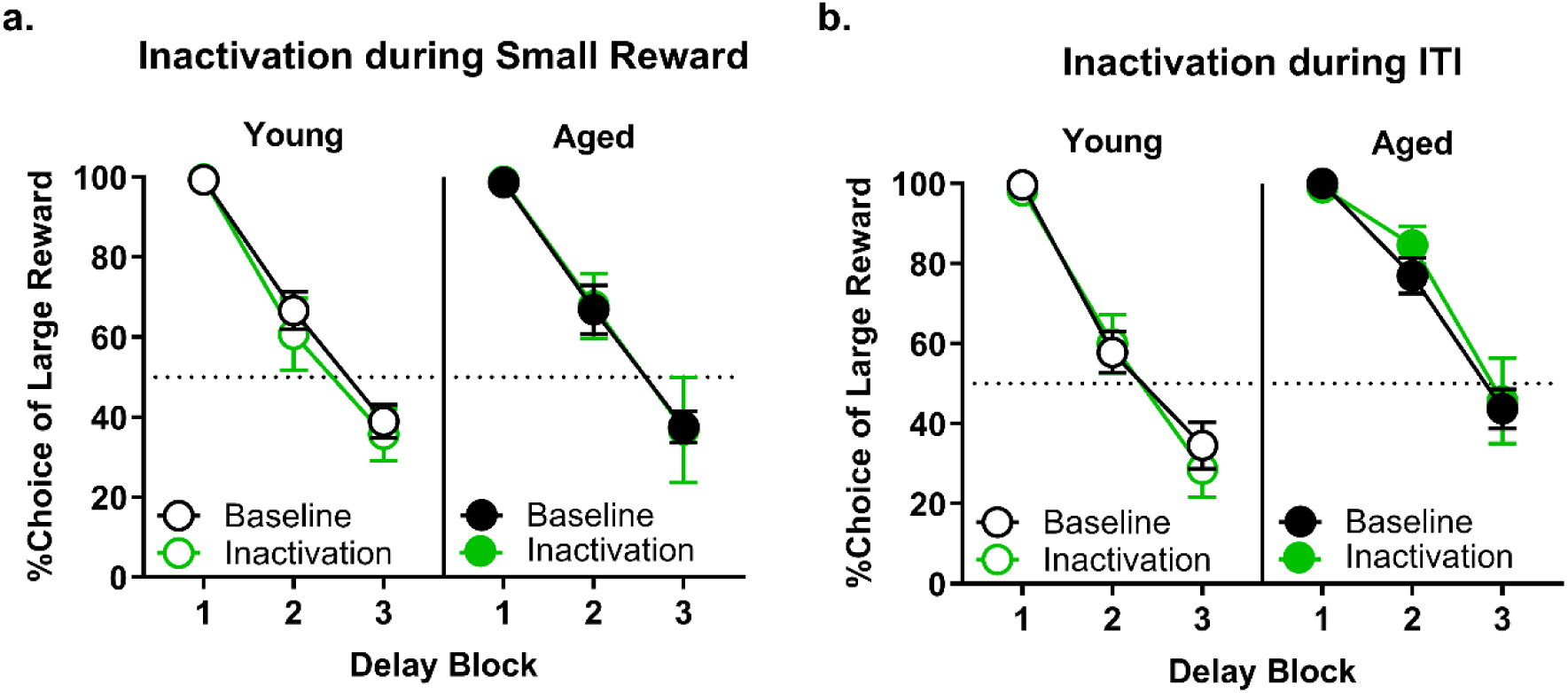
Effect of mPFC inactivation during the small reward and intertrial interval (ITI) epochs. **a)** Inactivation of the mPFC during the small reward epoch resulted in no change in choice performance in either young (n=12) or aged (n=6) rats. **b)** Inactivation of the mPFC during the intertrial interval (ITI) epoch resulted in no change in choice performance in either young (n=12) or aged (n=6) rats. Error bars represent standard error of the mean (SEM).

### Effects of mPFC inactivation during the intertrial interval

To confirm the temporal specificity of the mPFC inactivation effects, rats (n=12 young, n=6 aged) were tested while the mPFC was inactivated during the intertrial interval (ITI) (Figure 7b). Although the expected main effect of delay block was observed (F_(1.72,27.44)_=109.99, p<0.01, η_p_^2^=0.87, 1-β=1.00), mPFC inactivation during the ITI did not alter choice performance compared to baseline in young or aged rats (main effect of laser condition: F_(1,16)_=0.04, p=0.85; laser condition × age: F_(1,16)_=0.69, p=0.42; laser condition × delay block: F_(1.58,25.33)_=1.35, p=0.27; laser condition × age × delay block: F_(1.58,25.33)_=0.34, p=0.66). There was a main effect of age due to a difference in baseline performance (F_(1,16)_=4.35, p=0.05), but there was no interaction between age and delay block (F_(1.72,27.44)_=3.39, p=0.06).

#### Effects of light delivery into mPFC in rats with control virus (AAV5-CamkIIα-mCherry)

To control for non-specific effects of light delivery (e.g., changes in tissue temperature), the effects of light delivery in rats transduced with a control virus that did not contain the eNpHR3.0 gene were tested during behavioral epochs in which mPFC inactivation influenced choice behavior (i.e., deliberation: n=5 young and n=3 aged rats; delay: n=5 young and n=5 aged rats; and large reward: n=5 young and n=3 aged rats).

#### Effects of light delivery during the deliberation epoch

Light delivery during the deliberation epoch in rats transduced with a control virus had no effects on choice performance (Figure 8a). A three factor ANOVA (laser condition × age × delay block) indicated the expected main effect of delay block (F_(1.64,9.86)_=76.07, p<0.01, η_p_^2^=0.93, 1-β=1.00) but no significant main effects or interactions involving laser condition or age (main effect of laser condition: F_(1,6)_<0.01, p=0.94; main effect of age: F_(1,6)_=5.65, p=0.06; laser condition × age: F_(1,6)_<0.01, p=0.93; laser condition × delay block: F_(1.86,11.14)_=0.38, p=0.69; age × delay block: F_(1.64,9.86)_=2.00, p=0.19; laser condition × age × delay block: F_(1.86,11.14)_=0.57, p=0.57).

**Figure 8.**
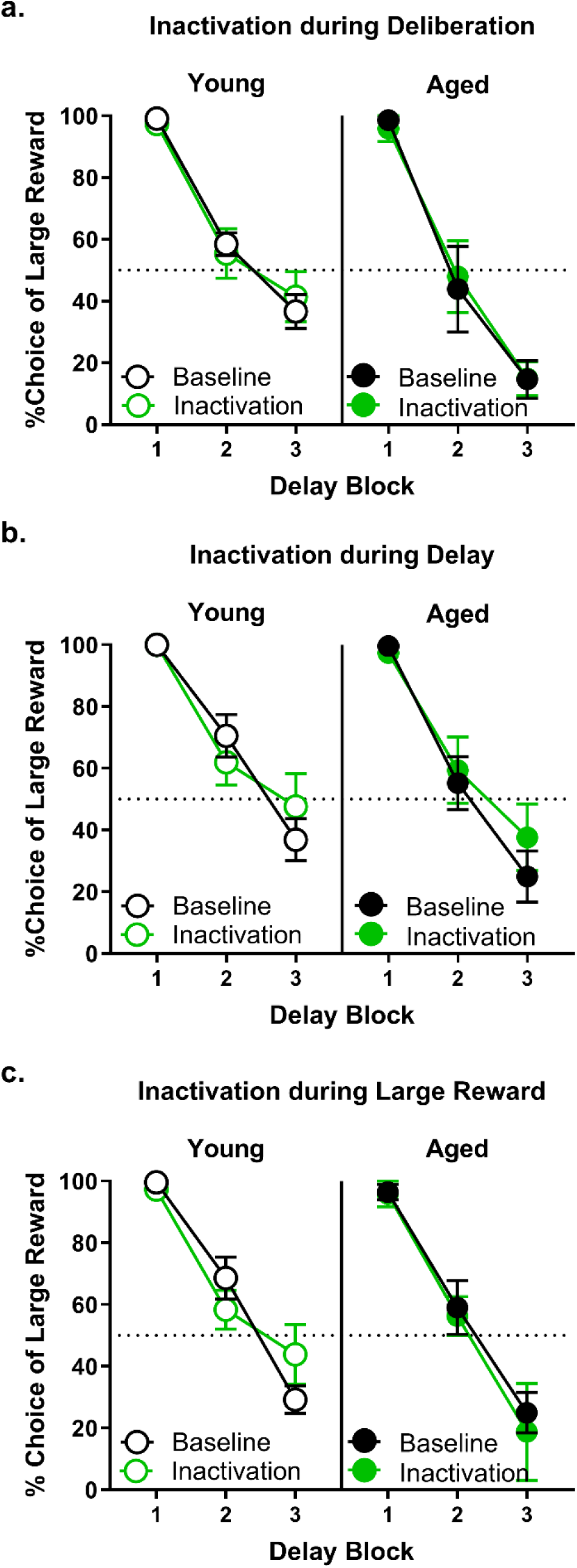
Effect of light delivery into mPFC in rats transduced with the control virus (AAV5-CamkIIα-mCherry). Light delivery into the mPFC in rats transduced with a control virus during. **a)** the deliberation, **b)** delay, and **c)** large reward epochs resulted in no change in choice performance in either young or aged rats (deliberation: n=5 young and n=3 aged rats; delay: n=5 young and n=5 aged rats; and large reward: n=5 young and n=3 aged rats). Error bars represent standard error of the mean (SEM).

#### Effects of light delivery during the delay reward epoch

Light delivery during the delay epoch in rats transduced with a control virus had no effects on choice performance (Figure 8b). A three factor ANOVA (laser condition × age × delay block) indicated the expected main effect of delay block (F_(1,8)_=30.88, p<0.01, η_p_^2^=0.79, 1-β=1.00) but no significant main effects or interactions involving laser condition or age (main effect of laser condition: F_(1,8)_=3.67, p=0.09; main effect of age: F_(1,8)_=0.87, p=0.38; laser condition × age: F_(1,8)_=2.19, p=0.18; laser condition × delay block: F_(1,8)_=2.70, p=0.14; age × delay block: F_(1,8)_=0.05, p=0.83; laser condition × age × delay block: F_(1,8)_=0.41, p=0.54).

#### Effects of light delivery during the large reward epoch

Light delivery during the large reward epoch in rats transduced with a control virus had no effects on choice performance (Figure 8c). A three factor ANOVA (laser condition × age × delay block) indicated the expected main effect of delay block (F_(1.95,11.68)_=81.90, p<0.01, η_p_^2^=0.93, 1-β=1.00) but no significant main effects or interactions involving laser condition or age (main effect of laser condition: F_(1,6)_=0.2, p=0.67; main effect of age: F_(1,6)_=1.61, p=0.25; laser condition × age: F_(1,6)_=0.48, p=0.51; laser condition × delay block: F_(1.23,7.41)_=1.20, p=0.32; age × delay block: F_(1.95,11.68)_=0.72, p=0.50; laser condition × age × delay block: F_(1.24,7.41)_=2.32, p=0.17).

## Discussion

Aging is accompanied by alterations in multiple forms of cost-benefit decision-making that are mediated by the prefrontal cortex (Samson, Venkatesh et al. 2015, Hernandez, Vetere et al. 2017, Lighthall 2020, Orsini, Pyon et al. 2023). Here we used an optogenetic approach in a rat model that recapitulates age-related shifts in intertemporal choices behavior observed in humans (Jimura, Myerson et al. 2011, Eppinger, Nystrom et al. 2012, Roesch, Bryden et al. 2012, Beas, Setlow et al. 2015, Hernandez, Vetere et al. 2017, Hernandez, Orsini et al. 2019, Hernandez, McQuail et al. 2022), to determine the contributions of mPFC to different stages of the decision-making process (deliberation between choice options, the delay prior to large reward delivery, and receipt of the two reward outcomes) and how these contributions differ in young and aged rats. The results show that mPFC inactivation produces both common and age-specific shifts in choice behavior across different stages of the decision process, highlighting potentially unique roles for the mPFC in regulating intertemporal choice across the lifespan.

### mPFC inactivation during deliberation

Inactivation of mPFC during deliberation increased preference for large, delayed over small immediate rewards (i.e., reduced impulsivity) in rats of both ages. This increase in choice of large, delayed rewards was evident after a prior choice of either a small or large reward, suggesting that it reflects an alteration in deliberative decision-making, rather than impaired behavioral flexibility. In addition, the fact that mPFC inactivation increased latencies to press the small reward lever is suggestive of a reduction in the relative valuation of the small reward (Hernandez, Vetere et al. 2017), and consistent with the pattern of choice behavior itself. The increase in large reward preference following mPFC inactivation during deliberation is similar to that observed following BLA inactivation during the same task epoch (Hernandez, Orsini et al. 2019), consistent with the strong reciprocal connections between the two brain regions as well as evidence that they act as a functional circuit mediating intertemporal choice and other forms of decision making (St Onge, Stopper et al. 2012). Considered together, these findings suggest that under normal (intact brain) conditions, activity in the mPFC-BLA circuit during deliberation biases subsequent choices toward smaller, more immediate rewards. In addition, the fact that inactivation of either structure during deliberation replicates the aged phenotype (greater preference for large, delayed over small, immediate rewards (Hernandez, Vetere et al. 2017)), could suggest that aging is accompanied by hypoactivity in this circuit during this phase of the decision process.

Two recent studies also investigated the effects of optogenetic mPFC manipulations during deliberation on intertemporal choice in young rats. In one, mPFC inactivation reduced preference for large, delayed rewards (White, Morningstar et al. 2024), whereas in the other, mPFC activation had no effect on delay preference but disrupted rats’ choice strategies (McLaughlin and Redish 2023). Notably, unlike the present study that employed a “fixed delay” design in which delays to large reward delivery remained constant within a block of trials, these prior studies employed designs in which either the small reward magnitude (White, Morningstar et al. 2024) or large reward delay (McLaughlin and Redish 2023) was adjusted within sessions as a function of rats’ choices. As these “adjusting” designs likely place greater demands on cognitive flexibility, it is likely that this factor accounts at least in part for the distinct effects of these manipulations.

### mPFC inactivation during the delay to large reward delivery

Similar to its effects during deliberation, inactivation of mPFC during delays prior to large reward delivery increased preference for the large reward. In contrast to its effects during deliberation, however, this effect was only evident in aged and not young rats. The reasons for this unique effect in aged rats are not clear, particularly given that neural representations of subjective value during intertemporal choices appear to be maintained across the lifespan (Seaman, Brooks et al. 2018). Although it is possible that young rats do not critically engage their mPFC during the delay period, this seems unlikely given extensive evidence that this brain region (and homologous PFC regions in non-human primates) is engaged in maintaining task-relevant information during delay periods in young adults during both intertemporal choice and related tasks (Rossi, Hayrapetyan et al. 2012, Arnsten and Jin 2014, Cao, Hong et al. 2023, McLaughlin and Redish 2023, White, Morningstar et al. 2024). One possibility is that representations maintained by mPFC activity that link responses on the large reward lever with anticipated outcomes during the delay period are simply more fragile in aged compared to young rats, and thus more readily disrupted by optogenetic inactivation. This interpretation is consistent with findings in non-human primates that neural activity encoding to-be-remembered information decays more rapidly in aged compared to young PFC (Wang, Gamo et al. 2011). A second (not mutually exclusive) possibility is that young rats are able to rapidly integrate delays into the value of the large reward (e.g., during the forced-choice trials that precede each block of free-choice trials), such that performance quickly becomes independent of mPFC activity during the delay period, whereas this process is slower in aged rats and thus more subject to disruption. Yet a third possibility is that young rats are able to use redundant neural systems to perceive and/or integrate reward delays that bypass mPFC activity, whereas aged rats do not have access to such redundancy.

Irrespective of the explanation, it is not necessarily clear why disrupting representations linking choices to outcomes prior to large reward delivery would enhance preference for this reward. It is possible, however, that disrupted representation or integration of reward delays renders aged rats less able to take such delays into account when calculating the relative values of the large vs. the small reward (i.e., delays would fail to discount the value of the large reward), rendering their choices more biased by reward magnitude alone (e.g., maintained by stimulus-response associative learning). Indeed, other studies show that mPFC (in young subjects) plays a critical role in some aspects of reward revaluation (Corbit and Balleine 2003, Ostlund and Balleine 2005), and that aging itself can impair some (though not all) forms of revaluation (Singh, Jones et al. 2011, Gray, Umapathy et al. 2018). Importantly, it is not likely the case that the different effects of mPFC inactivation during the delay in young and aged rats are driven by differences in halorhodopsin expression or function. Both the extent of expression and *ex vivo* function of halrhodopsin were comparable in young and aged rats. In addition, mPFC inactivation during deliberation was equally effective at modulating behavior at both ages, suggesting that differential effects of inactivation during delays resulted from age differences in brain function rather than opsin efficacy (see also (Hernandez, Orsini et al. 2019)). Moreover, although aging itself can cause shifts in time estimation that have the potential to contribute to altered intertemporal choice, altered perception of the delay duration is unlikely to be responsible for the inactivation-induced increase in preference for the large reward, as lesions or pharmacological inactivation of mPFC do not disrupt time estimation (Narayanan, Horst et al. 2006, Buhusi, Reyes et al. 2018).

In contrast to mPFC inactivation during the delay, BLA inactivation during the same epoch has no effects on choice behavior (Hernandez, Orsini et al. 2019). This pattern of results, in combination with those from inactivation of the same brain regions during deliberation, indicates that the involvement in intertemporal choice of the mPFC-BLA circuit is limited to only some phases of the decision-making process. More broadly, our prior findings are consistent with the general absence of impairing effects of BLA dysfunction in maintenance of (non-affective) information across delays (Roozendaal, McReynolds et al. 2004, Morgan, Terburg et al. 2012).

### mPFC inactivation during other epochs

Inactivation of mPFC during delivery of either the small or the large reward did not significantly alter choice behavior, although the interaction between age, laser condition, and block was near significant in the case of the large reward. In addition, there was no effect of mPFC inactivation during the inter-trial interval, nor were there effects of light delivery in control rats transduced with mCherry alone. These results indicate that the critical role of mPFC in intertemporal choice is limited to the deliberation and delay phases of the task, and that these results are not readily explained by non-specific effects of light delivery.

This pattern of results also contrasts with the effects of BLA inactivation during outcome delivery, which caused a reduction in choice of the large, delayed reward in young but not aged rats when inactivation took place during the small reward outcome (Hernandez, Orsini et al. 2019). Considered together, these data indicate that, despite both mPFC and BLA being critical for intertemporal choice, they play overlapping and distinct roles in different components of the decision process across the lifespan. Both structures appear to act in concert during deliberation to promote preference for immediate gratification. During delays prior to large reward delivery, the BLA does not seem to be engaged, whereas mPFC plays a unique role in aged rats in representing or integrating delays to modulate the subjective value of the large reward. Finally, during outcome evaluation, the mPFC does not seem to be critical at all, whereas the BLA is uniquely engaged in young rats in using the experience of immediate gratification to direct future behavior toward delayed (and larger) reward choices. Such distinct roles for the two structures are consistent with recent data showing that pharmacological activation of GABA(B) receptors in mPFC and BLA has both age-specific and directionally-distinct effects on intertemporal choice (Hernandez, McQuail et al. 2022).

### The mPFC in the context of the neural circuitry of intertemporal choice

Early work on neural mechanisms of intertemporal choice in rodents showed mixed effects of lesions or other long-lasting manipulations of mPFC, with both decreases and no effects on preference for large, delayed rewards reported (e.g., (Cardinal, Pennicott et al. 2001, Churchwell, Morris et al. 2009, Loos, Pattij et al. 2010, Feja and Koch 2014, Sonntag, Brenhouse et al. 2014, Yates, Perry et al. 2014, Déziel and Tasker 2017)), despite good evidence linking mPFC monoamine signaling to intertemporal choice (Winstanley, Theobald et al. 2006, Loos, Pattij et al. 2010, Simon, Beas et al. 2013). More recently, electrophysiological recording studies have shown that mPFC single-unit activity encodes upcoming choices and anticipated rewards in intertemporal choice tasks (Sackett, Moschak et al. 2019, Cao, Hong et al. 2023), as well as shifts in preference across multiple choice trials (White, Morningstar et al. 2024). It is likely that this different task-related activity reflects distinct inputs and outputs of mPFC to other brain structures critical for intertemporal choice (most notably BLA, but also OFC, NAc, and vHPC), as well as dopaminergic and serotonergic innervation (Winstanley, Theobald et al. 2006, Yates, Perry et al. 2014, Bailey, Simpson et al. 2016, Fobbs and Mizumori 2017). With few exceptions, however (Wenzel, Zlebnik et al. 2023), and in contrast to other forms of cost-benefit decision making (St Onge, Stopper et al. 2012, Jenni, Larkin et al. 2017), circuit-based approaches to investigating the contributions of mPFC to intertemporal choice have been understudied. As such, understanding of the temporally- and circuit-specific contributions of mPFC activity to intertemporal choice, and the flow of information among nodes within mPFC-containing neural circuitry, remains limited. An additional limitation of the present work is that only males were used. Although data in humans suggest equivalent increases in preference for delayed gratification in men and women in aging (and, at least in young rats, mPFC encoding of intertemporal choice-related variables is similar in males and females; (Sackett, Moschak et al. 2019)), to our knowledge, intertemporal choice has not been characterized in aged female rodents, and remains an important direction of future research.

## Acknowledgements.

We thank Vicky S. Kelly and Bonnie I. McLaurin for assistance with surgeries and behavioral testing.

## Abbreviations

(mPFC): medial prefrontal cortex;
(FBN): Fischer 344 x Brown Norway hybrid;
(ITI): intertrial interval

